# TCF7L2 plays a complex role in human adipose progenitor biology which may contribute to genetic susceptibility to type 2 diabetes

**DOI:** 10.1101/854661

**Authors:** Manu Verma, Nellie Y. Loh, Senthil K. Vasan, Andrea D. van Dam, Marijana Todorčević, Matt J. Neville, Fredrik Karpe, Constantinos Christodoulides

## Abstract

Non-coding genetic variation at *TCF7L2* is the strongest genetic determinant of type 2 diabetes (T2D) risk in humans. TCF7L2 encodes a transcription factor mediating the nuclear effects of WNT signalling in adipose tissue (AT). Here we mapped the expression of *TCF7L2* in human AT and investigated its role in adipose progenitor (AP) biology. APs exhibited the highest *TCF7L2* mRNA abundance compared to mature adipocytes and adipose-derived endothelial cells. Obesity was associated with reduced *TCF7L2* transcript levels in subcutaneous abdominal AT but increased expression in APs. In functional studies, TCF7L2 knockdown (KD) in APs led to dose-dependent activation of WNT/β-catenin signaling, impaired proliferation and dose-dependent effects on adipogenesis. Whilst partial KD enhanced adipocyte differentiation, complete KD impaired lipid accumulation and adipogenic gene expression. Overexpression of TCF7L2 accelerated adipogenesis. Transcriptome-wide profiling revealed that TCF7L2 can modulate multiple aspects of AP biology including extracellular matrix secretion, immune signalling and apoptosis. The T2D-risk allele at rs7903146 was associated with reduced AP *TCF7L2* expression and enhanced AT insulin sensitivity. Our study highlights a complex role for TCF7L2 in AP biology and suggests that in addition to regulating pancreatic insulin secretion, genetic variation at *TCF7L2* may also influence T2D risk by modulating AP function.

## INTRODUCTION

Adipose tissue (AT) plays a central role in the regulation of whole-body energy and glucose homoeostasis. Firstly, it provides a safe depot for the storage of excess calories in the form of triglycerides, thereby protecting extra-adipose tissues from lipotoxicity (1). Secondly, it releases energy as free fatty acids during periods of energy demand such as fasting and exercise. Finally, through the secretion of peptide hormones such as leptin and adiponectin it directly regulates systemic energy balance and insulin sensitivity (2). AT expands through an increase in adipocyte number (hyperplasia) and/or size (hypertrophy). Hyperplastic AT growth is mediated *via* generation of new adipocytes from adipose progenitors (APs) and is associated with enhanced systemic insulin sensitivity and an improved glucose and lipid profile. In contrast, adipocyte hypertrophy results in AT inflammation and fibrosis which lead to the development of insulin resistance (3,4).

WNTs are a family of secreted growth factors which function in autocrine and paracrine fashions to regulate stem cell and AP biology (5–7). WNTs signal through frizzled (FZD) receptors to activate multiple downstream signalling cascades. In the best characterized, canonical pathway, WNT binding to FZDs leads to nuclear accumulation of the transcriptional co-activator β-catenin which in conjunction with TCF/LEF transcription factors activates WNT target gene expression. In the absence of WNTs, TCF/LEF proteins are bound to members of the Groucho/TLE family of transcriptional co-repressors and suppress WNT/β-catenin signalling. There are four TCF/LEF family members in mammals; TCF7, TCF7L1, TCF7L2 and LEF1 (8). Of these, TCF7L2 is the most highly expressed in AT (9).

Work from the MacDougald laboratory first highlighted a potential role for TCF7L2 in the regulation of AT function (10). A dominant negative TCF7L2 mutant lacking its β-catenin binding domain was shown to cause spontaneous adipogenesis of 3T3-L1 cells and trans-differentiation of C2C12 myoblasts into adipocytes. More recently, the *in vitro* role of TCF7L2 in the regulation of adipogenesis was revisited with conflicting results. Whilst Chen *et al* showed that TCF7L2 knockdown (KD) in 3T3-L1 preadipocytes leads to impaired adipocyte differentiation (11) in another study inducible TCF7L2 deletion in immortalised inguinal mouse APs was associated with enhanced adipogenesis (12). *Ex vivo* adipose expression of *Tcf7l2* was suppressed by high-fat diet (HFD)-induced and genetic obesity (12,13). *In vivo*, homozygous global *Tcf7l2* null mice die perinatally whereas heterozygous null animals are lean with enhanced glucose tolerance and insulin sensitivity compared to wild-type controls both on a chow and following a HFD (14). Conversely, transgenic mice over-expressing *Tcf7l2* displayed HFD-induced glucose intolerance (14). Contrasting these findings, targeted deletion of *Tcf7l2* in mature adipocytes (mADs) resulted in enhanced adiposity, glucose intolerance and hyperinsulinaemia (11,12). In humans, *TCF7L2* expression was decreased in the SC abdominal (hereafter referred to as abdominal) AT of subjects with impaired glucose tolerance (11) and reduced systemic insulin sensitivity (15). Notably, non-coding genetic variation at the *TCF7L2* locus was demonstrated to be the strongest genetic determinant of type 2 diabetes (T2D) risk in humans (16,17). The same single nucleotide variation (SNV), rs7903146, was also associated with fat distribution (18). Prompted by these findings we investigated the role of TCF7L2 in human AT function.

## RESEARCH DESIGN AND METHODS

### Study population

Study subjects were recruited from the Oxford Biobank (www.oxfordbiobank.org.uk), a population-based cohort of healthy 30-50-year-old volunteers (19). Paired abdominal and visceral biopsies were collected from patients undergoing elective surgery as a part of the MolSURG study. Plasma biochemistry (19) and adipocyte sizing (20) were undertaken as described. All studies were approved by the Oxfordshire Clinical Research Ethics Committee, and all volunteers gave written, informed consent.

### Cell culture

Primary APs (derived from AT biopsies) or AP lines were cultured and differentiated as described (21,22). Primary endothelial cells were isolated using a CD31 MicroBead Kit (Miltenyi Biotec). Quantification of intracellular lipid was undertaken using AdipoRed lipid stain (Lonza) and multi-well plate reader (PerSeptive Biosystems, Perkin Elmer).

### Generation of de-differentiated fat (DFAT) cells

DFAT cells were generated by selection and de-differentiation of lipid-laden, *in vitro* differentiated immortalised APs (23) with modifications (See supplemental information).

### Lentiviral constructs and generation of stable AP lines

TCF7L2 (sh843, TRCN0000262843; sh897, TRCN0000061897) and control (scrambled) shRNA plasmid vectors were purchased (Sigma-Aldrich). The TOPflash reporter vector (24) was a gift from Roel Nusse (Addgene #24307). Lentiviral particles were produced in HEK293 cells using MISSION® (Sigma-Aldrich) packaging mix. Stable AP lines were generated by transduction of cells with lentiviral particles and selected using (2μg/ml) puromycin.

### Doxycycline-inducible AP lines

The *TCF7L2* sequence (from TCF4E pcDNA3, a kind gift from Frank McCormick, Addgene #32738) (25) was cloned into the tet-pLKO-puro doxycycline-inducible expression lentiviral vector (gift of Dmitri Wiederschain, Addgene #21915) (26). Stable doxycycline-inducible AP lines were generated by transduction of cells with lentiviral particles and selected using (2μg/ml) puromycin.

### Proliferation assays

Equal number of APs were seeded, trypsinised and counted using a Cellometer Auto T4 (Nexcelom Bioscience) every 96 hours or using CyQUANT™ cell proliferation assays. Doubling time was calculated using the formula: *T_d_* = (*t_2_* − *t_1_*) x [log (2) ÷ log (*q_2_* ÷ *q_1_*)], where *t* = time (days), *q* = cell number.

### Luciferase assays

To asses cis-regulatory activity, a 150 base pair genomic sequence centred around rs7903146 was cloned into a luciferase reporter vector (pGL4.23[luc2/minP], Promega) and co-transfected with pRL-SV40 (Promega) into HEK293 (Lipofectamine 2000) or DFAT cells (Neon Transfection System). 48h post transfection, luciferase activity was assessed using the Dual-Luciferase® Reporter Assay System (Promega) on Veritas Microplate Luminometer (Turner Biosystems).

### TOPflash reporter assays

β-catenin transcriptional activity in TCF7L2 KD or overexpressing DFAT cells was determined as described (21).

### Real time-PCR and western blots

qRT-PCR and western blotting were performed using TaqMan assays and standard protocols (see supplemental information).

### RNA sequencing, pathway enrichment and transcription factor binding-site motif analysis

RNA sequencing (RNA-Seq) was performed in scrambled control, sh897 and sh843 TCF7L2-KD DFAT abdominal APs (see supplemental information). Differentially regulated genes with a false discovery rate <0.05 and an absolute fold-change >1.5, were selected for further analysis. Pathway enrichment and transcription factor binding-site motif analysis were undertaken using Metascape (27) and iRegulon (28), respectively.

Statistical analysis: Statistical analysis was performed using R, SPSS, STATA or GraphPad. For parametric data, Pearson’s correlation, two-tailed student’s test with Welch’s correction (where appropriate) for 2 groups, or one- or two-way ANOVA followed by appropriate *post hoc* tests for multiple groups were used. For non-parametric data, Spearman’s correlation, two-tailed Mann-Whitney or Kruskal-Wallis tests were used. Smoothened splines using generalized additive models were used to investigate the relationship between Adipo-IR and BMI in different rs7903146 genotype carriers. p-values were corrected for multiple comparisons or age, BMI and sex (where appropriate) and p <0.05 was considered significant.

## RESULTS

### TCF7L2 expression is highest in APs

To elucidate the role of TCF7L2 in human AT biology we mapped its expression profile in AT. In biopsy samples from 30 lean and 30 obese individuals, *TCF7L2* mRNA abundance tended to be higher in abdominal *versus* gluteal fat (Fig. 1A). Compared with obese subjects, lean individuals had higher *TCF7L2* expression in abdominal AT. *TCF7L2* transcript levels were similar in the abdominal and visceral AT depots (Fig. 1B). In fractionated AT from over 100 healthy volunteers, *TCF7L2* mRNA levels were higher in APs compared with mADs from both the abdominal and gluteal fat depots (Fig. 1C). Furthermore, in a small sample group (n=5-6), *TCF7L2* expression was higher in APs *versus* adipose-derived endothelial cells (Fig. 1D). Finally, *TCF7L2* mRNA abundance in abdominal APs correlated positively with body mass index (BMI) whilst an opposite trend was detected in abdominal mADs (Fig. 1E, F). No associations between BMI and *TCF7L2* expression in gluteal adipose cell fractions were detected (Fig. S1). Based on these data we focused our subsequent studies on deciphering the function of TCF7L2 in APs.

**Fig. 1.**
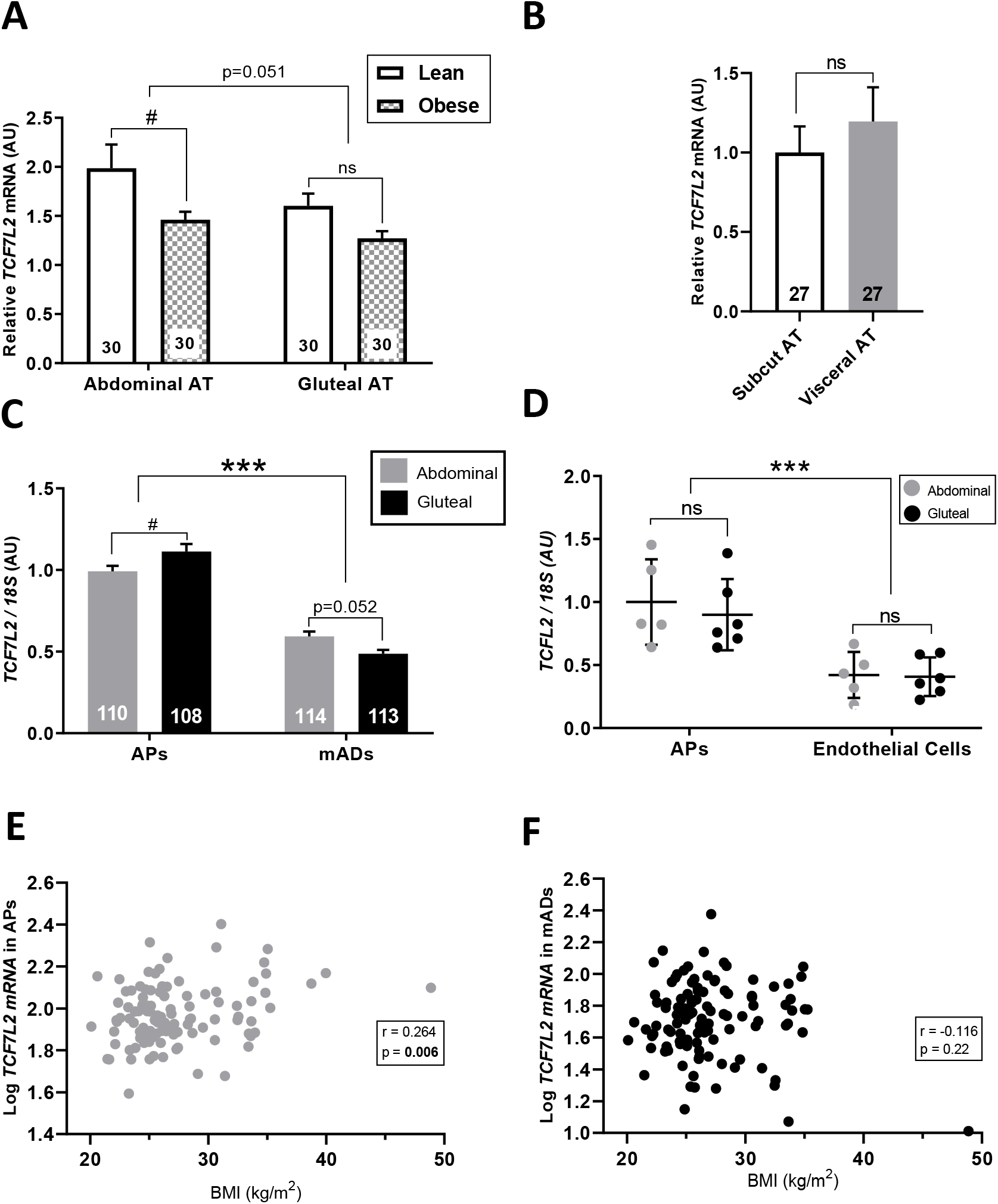
*Ex vivo TCF7L2* expression in human adipose depots and adipose cell fractions. *TCF7L2* expression in paired samples of (A) subcutaneous abdominal and gluteal adipose tissue (AT) biopsies from lean and obese subjects (n=30 [15F] / group; Lean – Age 43.9±4.7 years, BMI 22.0±1.1 kg/m^2^; Obese – Age 43.9±3.6 years, BMI 34.5±2.4 kg/m^2^), (B) abdominal subcutaneous and visceral AT biopsies (n=27 [16F] / group; Age - 59.1±11.8 years, BMI - 28.2±7.2 kg/m^2^), (C) cultured APs and mature adipocytes (mADs) from subcutaneous abdominal and gluteal AT biopsies (n=108-114 [59F] / group; Age - 45.3±8.7 years, BMI - 27.2±4.5 kg/m^2^), and (D) cultured APs and adipose-derived endothelial cells from subcutaneous abdominal and gluteal AT biopsies (n=5-6 [5-6F] / group; Age - 51.9±7.9 years, BMI - 28.8±4.0 kg/m^2^). Errror bars are mean ± SD. (E, F) Correlations between *TCF7L2* expression in cultured APs and mADs from subcutaneous abdominal AT biopsies and donor BMI (n=110-113 [59F] / group; Age - 45.3±8.7 years, BMI - 27.2±4.5 kg/m^2^). qRT-PCR data were normalized to geometric mean of (A) *PPIA, PGK1, PSMB6*, and *IPO8*, (B) *PPIA* and *PGK1*, or (C-F) to 18S rRNA levels. ***p<0.001; #p<0.05 (adjusted for multiple comparisons). Histograms are means ± SEM. Age and BMI data are means ± SD.

### TCF7L2 dose-dependently modulates AP differentiation

We investigated the role of TCF7L2 in AP biology using de-differentiated fat (DFAT) cells generated from immortalised abdominal adipocytes (22). These retain their depot-specific gene expression signature (Fig. S2) and have a higher adipogenic capacity than primary and immortalised APs. Stable KD of TCF7L2 in these cells was achieved with two shRNAs targeting the universal exon 9 of *TCF7L2* (Fig. 2A, B). The first shRNA (sh897) led to 30% KD of TCF7L2 at mRNA level and 68% KD at protein level. The second, more efficient shRNA (sh843), led to 70% reduction in *TCF7L2* expression and near complete TCF7L2 protein KD. KD of TCF7L2 led to impaired AP proliferation (Fig. 2C). No differences in the doubling time of moderate-*versus* high-efficiency TCF7L2-KD cells were detected. Partial TCF7L2-KD was also associated with augmented adipocyte differentiation as ascertained by increased lipid accumulation and enhanced adipogenic gene expression (Fig. 2D - H). In sharp contrast, high-efficiency KD of TCF7L2 resulted in impaired adipogenesis. These findings were confirmed in primary abdominal TCF7L2-KD cells from two subjects. In these latter experiments the anti-adipogenic effect of sh843 was more subtle despite identical TCF7L2-KD efficiency in primary and DFAT cells (Fig. 2I, J). In complimentary experiments we also investigated the effects of inducible *TCF7L2* over-expression on AP functional characteristics using a Tet-On system (Fig. 3). The doxycycline (DOX) dose was selected to induce approximately 2- and 5-fold induction in *TCF7L2* expression compared to vehicle treated *TCF7L2* overexpressing APs which corresponded to an equivalent increase in protein production (and 2.6 and 6.6-fold higher protein production *versus* vehicle treated empty vector controls) (Fig. 3A, B). AP cells were grown in DOX-free media and *TCF7L2* expression was induced upon plating for proliferation and differentiation experiments and throughout thereafter. Ectopic expression of *TCF7L2* did not influence AP proliferation (Fig. 3C). Low-dose *TCF7L2* overexpression similarly did not affect adipocyte differentiation (Fig. S3) although, the DOX-induced increase in *TCF7L2* expression was not sustained throughout adipogenesis (Fig. S4). On the other hand, 5-fold increase in TCF7L2 protein production accelerated adipocyte differentiation (Fig. S5). By the end of the differentiation time course however, *TCF7L2* over-expressing adipocytes displayed only a subtle increase in lipid accumulation concomitant with increased *CEBPA* expression *versus* vehicle treated controls (Fig. 3D-H). We conclude that, *in vitro* TCF7L2 has dose-dependent actions on adipogenesis.

**Fig. 2.**
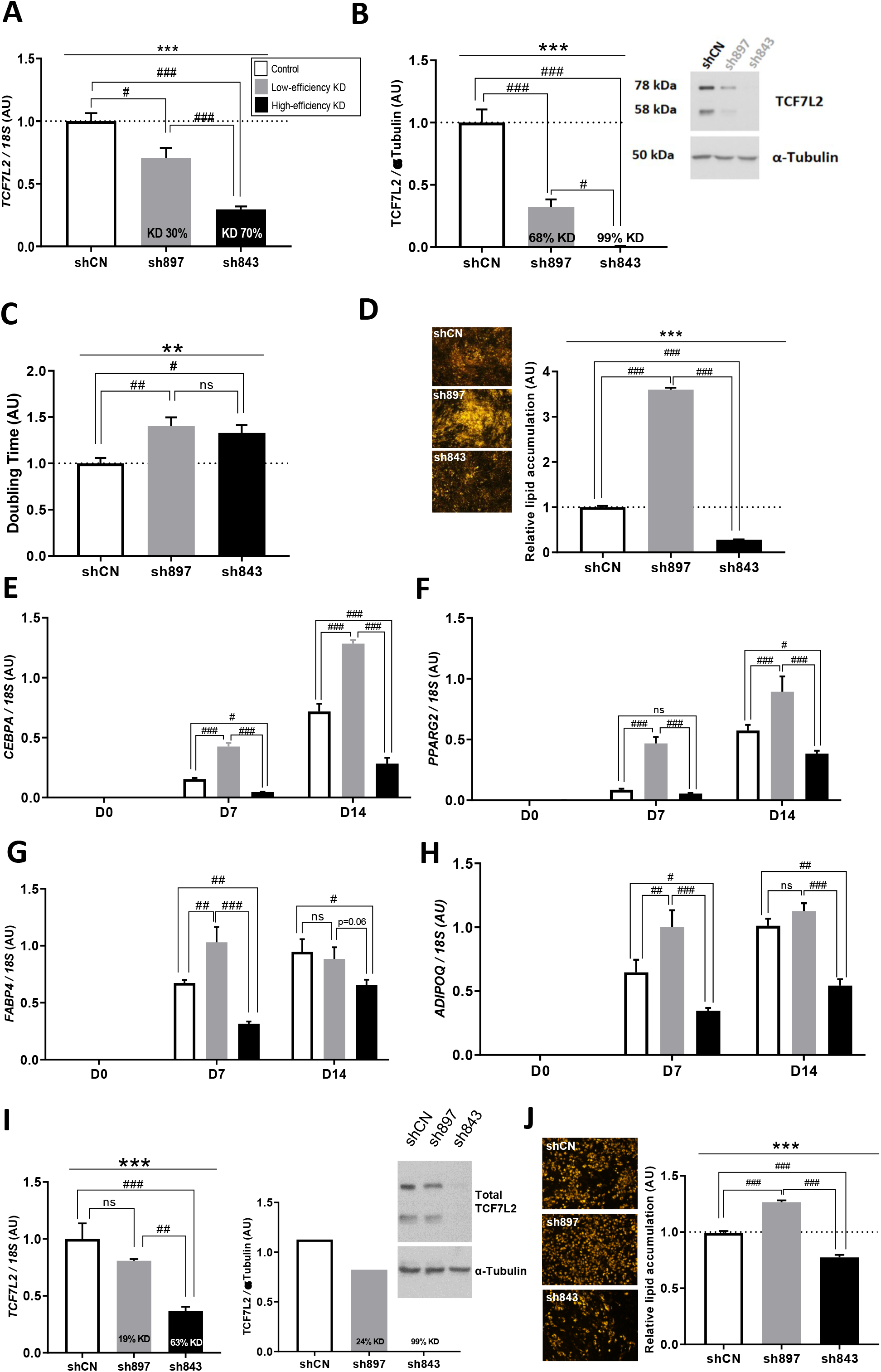
TCF7L2 KD in abdominal APs impairs proliferation and dose-dependently regulates differentiation. **(A-H)** TCF7L2 KD in DFAT abdominal APs. shCN = scrambled control, sh897 = moderate and sh843 = high TCF7L2 KD DFAT abdominal APs. TCF7L2 KD was confirmed by (A) qRT-PCR and (B) western blot in DFAT abdominal APs. (C) Doubling time of control, sh897 and sh843 TCF7L2 KD APs. (D) Representative micrographs of control, sh897 and sh843 APs at day 14 of adipogenic differentiation. The histogram shows the relative lipid accumulation (as a marker for differentiation) assessed by AdipoRed lipid stain (n=24 wells/group), (E-H) Relative mRNA levels of adipogenic genes *CEBPA, PPARG2, FABP4* and *ADIPOQ* at baseline (day 0), day 7 and day 14 of adipogenic differentiation. **(I-J)** TCF7L2 KD in human primary abdominal APs. (I) Confirmation of TCF7L2 KD by qRT-PCR and western blot in human primary abdominal APs (western blots from one experiment). (J) Representative micrographs of control, moderate (sh897) and high (sh843) TCF7L2 KD human primary abdominal APs at day 14 of adipogenic differentiation. The histogram shows the relative lipid accumulation, assessed by AdipoRed (n=16 wells/group). qRT-PCR data were normalized to 18S rRNA levels. Histograms are means ± SEM. Data obtained from 3 independent experiments. ***p<0.001, **p<0.01; ###p<0.001, ##p<0.01, #p<0.05 (adjusted for multiple comparisons). α-tubulin was used as a loading control for western blots.

**Fig. 3.**
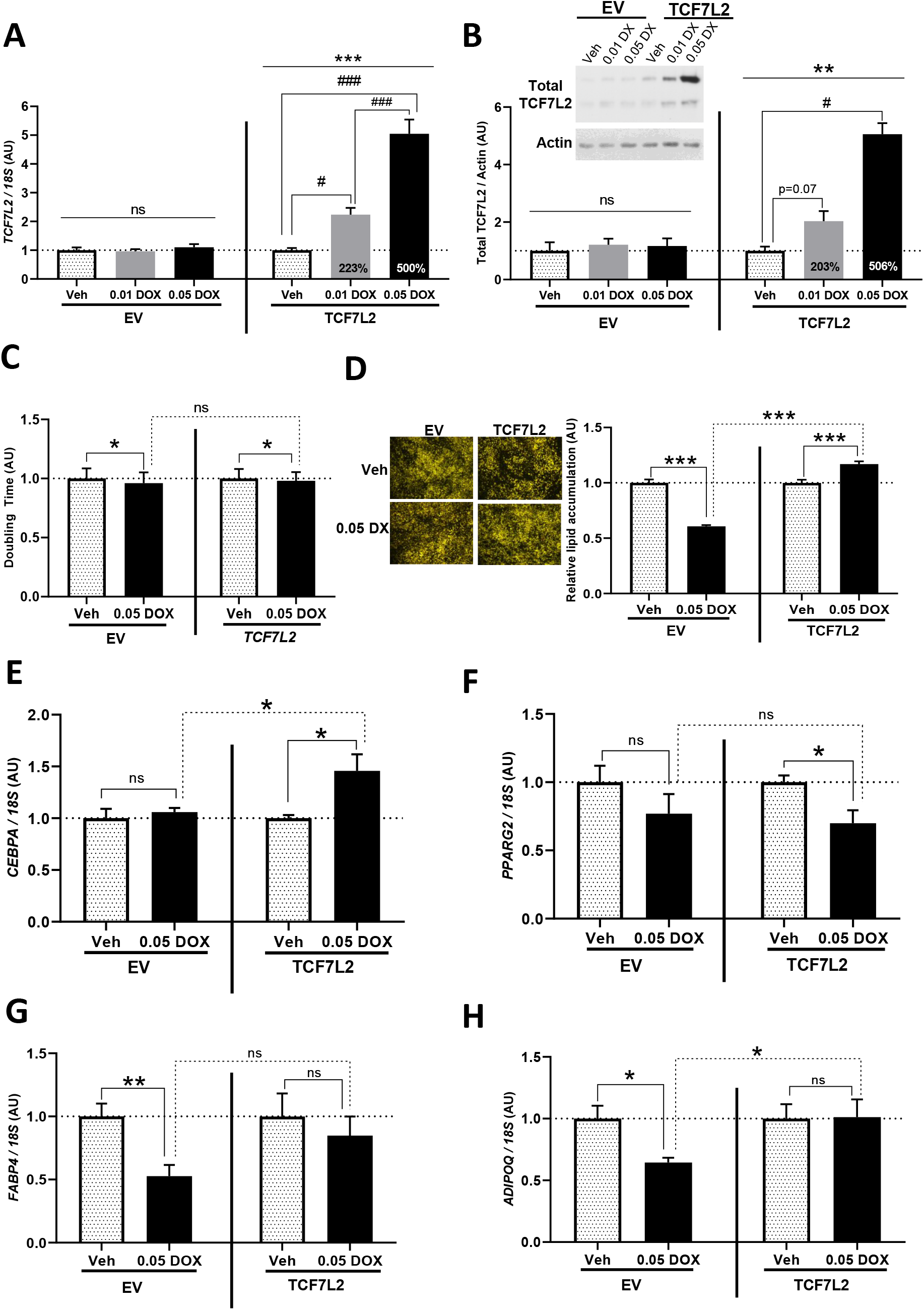
Doxycycline-induced TCF7L2 overexpression in DFAT abdominal APs regulates adipogenesis. DFAT abdominal APs, stably transduced with the empty vector (EV) or TCF7L2 overexpression vector (TCF7L2), were cultured in the presence of vehicle (Veh) or DOX (final concentration: 0.01 μg/ml or 0.05 μg/ml) to induce TCF7L2 overexpression. TCF7L2 overexpression was confirmed by (A) qRT-PCR and (B) western blot in DFAT abdominal APs. (C) Doubling time of DFAT abdominal APs. (D) Representative micrographs of DFAT abdominal APs at day 14 of adipogenic differentiation. The histogram shows the relative lipid accumulation, assessed by AdipoRed lipid stain (n=24 wells/group). (E-H) Relative mRNA levels of adipogenic genes *CEBPA, PPARG2, FABP4* and *ADIPOQ* at day 14 of adipogenic differentiation. qRT-PCR data were normalized to 18S rRNA levels. Histograms are means ± SEM and expressed relative to vehicle-treated cells (arbitrarily set to 1). Data obtained from 3 independent experiments. ***p<0.001, **p<0.01, *p<0.05; ###p<0.001, #p<0.05 (adjusted for multiple comparisons). Actin was used as western blot loading control.

### TCF7L2 dose-dependently modulates WNT/β-catenin signalling

We next determined whether the effects of TCF7L2 on AP function were driven by changes in WNT/β-catenin signalling (Fig. 4). Partial KD of TCF7L2 did not result in altered expression of the universal WNT target gene *AXIN2* (Fig. 4A). In contrast, (near) complete TCF7L2-KD was associated with increased *AXIN2* levels. Using TCF7L2-KD cells stably expressing the TOPflash promoter reporter construct which monitors endogenous β-catenin transcriptional activity we confirmed that complete depletion of TCF7L2 was associated with robust stimulation of canonical WNT signalling whilst weak activation was also seen following partial KD both in the absence and presence of WNT3A (Fig. 4B). Interestingly, despite increased β-catenin transcriptional activity, active β-catenin protein levels were decreased in both sh897 and sh843 cells (Fig. 4C). Given the paradoxical activation of WNT/β-catenin signalling in TCF7L2-KD APs we examined the expression of other TCF/LEF family members in these cells (Fig. 4D & Fig. S6A). Only *TCF7* and *TCF7L1* were expressed in abdominal APs. Notably, TCF7L2-KD was associated with a dose-dependent increase in *TCF7* mRNA abundance in both DFAT and primary abdominal APs (Fig. 4D). In gain-of-function experiments, 48-hour induction of 2- and 5-fold TCF7L2 production did not influence AP *AXIN2* mRNA levels (Fig. 4E). In TOPflash promoter assays, TCF7L2 over-expression led to inhibition of WNT/β-catenin signalling (Fig. 4F, G). However, DOX treatment also suppressed TOPflash activity in empty vector control cells. Over expression of TCF7L2 did not influence active β-catenin protein levels (Fig. 4I). Collectively these findings suggest that TCF7L2 functions to antagonize the transcriptional activity of β-catenin in abdominal APs.

**Fig. 4.**
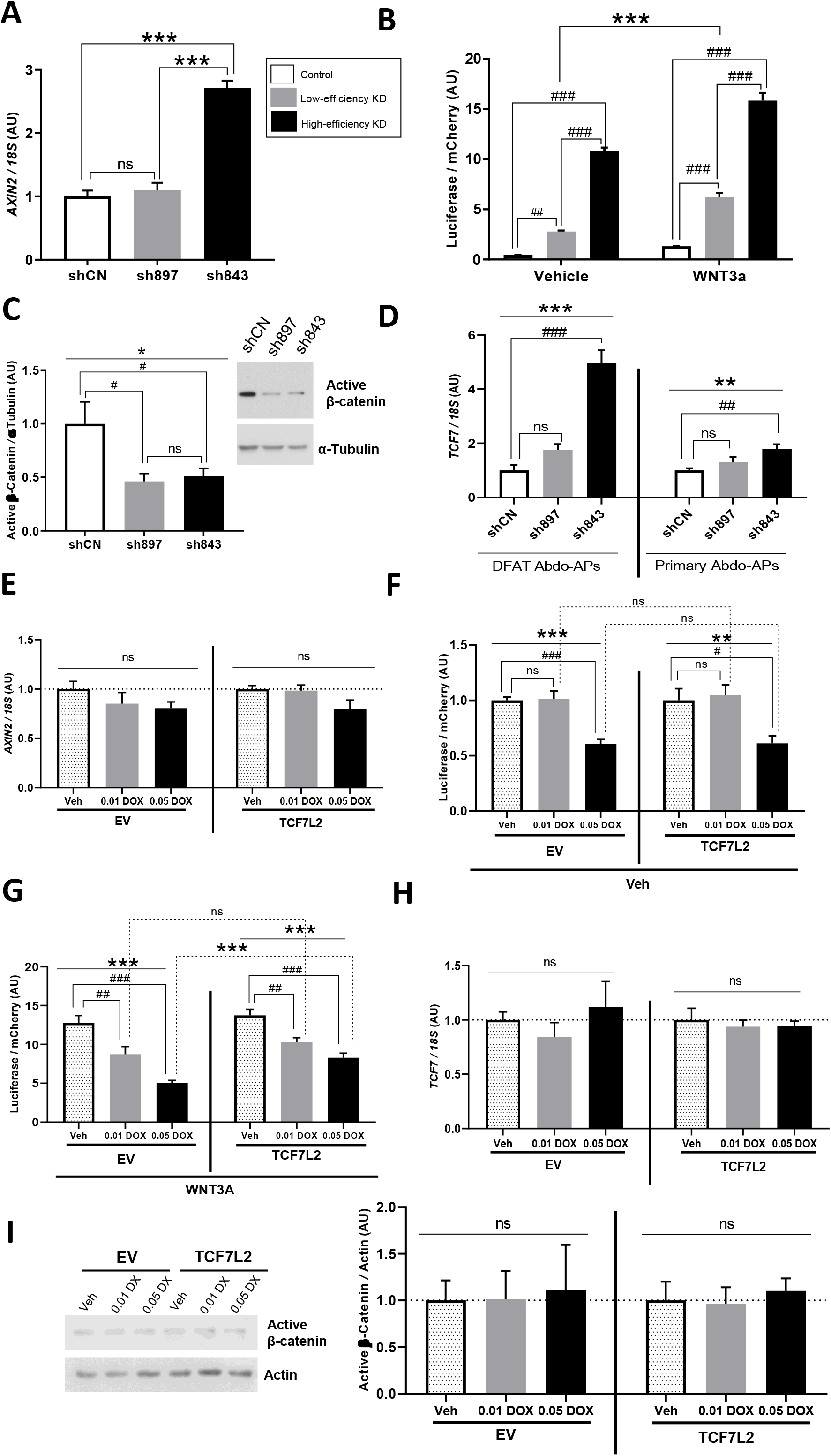
TCF7L2 dose-dependently modulates WNT/β-catenin signalling. (A-D) Effects of TCF7L2 KD on: (A) *AXIN2* mRNA levels, (B) TOPFlash promoter activity following 6h treatment with vehicle (Veh) or 50ng/ml WNT3A (n=12 wells/group), (C) active β-catenin protein levels, and (D) *TCF7* mRNA levels. shCN = scrambled control, sh897 = moderate and sh843 = high TCF7L2 KD DFAT abdominal APs. **(E-I)** Effects of TCF7L2 overexpression on: (E) *AXIN2* mRNA levels, (F and G) TOPFlash promoter activity following 20h treatment with (F) vehicle (Veh) or (G) 50ng/ml WNT3A (n=12 wells/group), (H) *TCF7* mRNA levels, and (I) active β-catenin protein levels. DFAT abdominal APs expressing either the empty vector (EV) or TCF7L2 overexpression vector (TCF7L2) were cultured in the presence of vehicle (Veh) or DOX (final concentration of 0.01 μg/ml or 0.05 μg/ml). Data expressed relative to vehicle treatment (arbitrarily set to 1) for EV or TCF7L2 overexpressing DFAT abdominal AP line, respectively. Actin was used as a loading control for western blots. qRT-PCR data were normalized to 18S rRNA levels. Histograms are means ± SEM. Data obtained from 3 independent experiments. ***p<0.001, **p<0.01, *p<0.05; ###p<0.001, ##p<0.01, #p<0.05 (adjusted for multiple comparisons).

### Transcriptome-wide profiling reveals that TCF7L2 regulates multiple aspects of AP biology

To identify the genes and biological processes regulated by TCF7L2 in APs we undertook transcriptomic analyses of TCF7L2-KD cells using RNA-Seq (Fig. 5A). High-efficiency TCF7L2-KD altered the expression of 2,806 genes (18% of all expressed genes) whilst partial KD resulted in a more modest change in the transcriptome with 1,024 genes differentially regulated (FDR < 5%, absolute fold-change > 1.5) (Fig. 5B). Gene set enrichment analysis revealed that the cluster of genes suppressed in complete TCF7L2-KD cells was enriched for pathways and processes involved in cell cycle, response to hypoxia, p53 signalling and ribosome biogenesis (Fig. 5E). Interestingly, cell cycle and ribosome biogenesis are two key pathways that are transcriptionally positively regulated by canonical WNT signalling (29). Consistent with these findings, the expression of many well-established WNT/β-catenin target genes was reduced in high-efficiency TCF7L2-KD APs (Fig. S7). The cluster of upregulated genes in high-efficiency KD cells was enriched for genes involved in interferon signalling and extracellular matrix (ECM) organization. In moderate-efficiency TCF7L2-KD cells the downregulated gene set was also enriched for pathways and processes involved in response to hypoxia and p53 signalling (Fig. 5F). Additionally, we detected enrichment for circadian regulation of gene expression. The upregulated gene cluster in sh897 cells was enriched for genes involved in ECM organization and immune and inflammatory processes. We also performed transcription factor-binding site motif analysis on the promoters of genes differentially expressed in TCF7L2-KD APs (Fig. 5E, F). The promoters of genes suppressed following both complete and partial TCF7L2-KD were enriched for binding sites of E2F4 and FOXM1 which regulate cell proliferation. Genes whose expression was decreased in high-efficiency TCF7L2-KD cells were also enriched for TFDP1, TFDP3 and KLF4 binding sites in their promoters. TFDP family members co-operatively regulate cell cycle genes with members of the E2F family whilst KLF4 suppresses p53 expression and is a β-catenin target gene. The promoters of genes suppressed in partial TCF7L2-KD cells were also enriched for HIF1A binding sites. Genes, whose expression was increased in both sh897 and sh843 cells, exhibited enrichment for binding sites of transcription factors involved in cytokine and particularly, interferon signalling in their promoters namely, IRF1, IRF7, STAT1, and STAT2. Finally, real-time PCR revealed good concordance between changes in the transcriptome of DFAT and primary APs following moderate-but not after high-efficiency TCF7L2-KD (Fig. 5G, H).

**Fig. 5.**
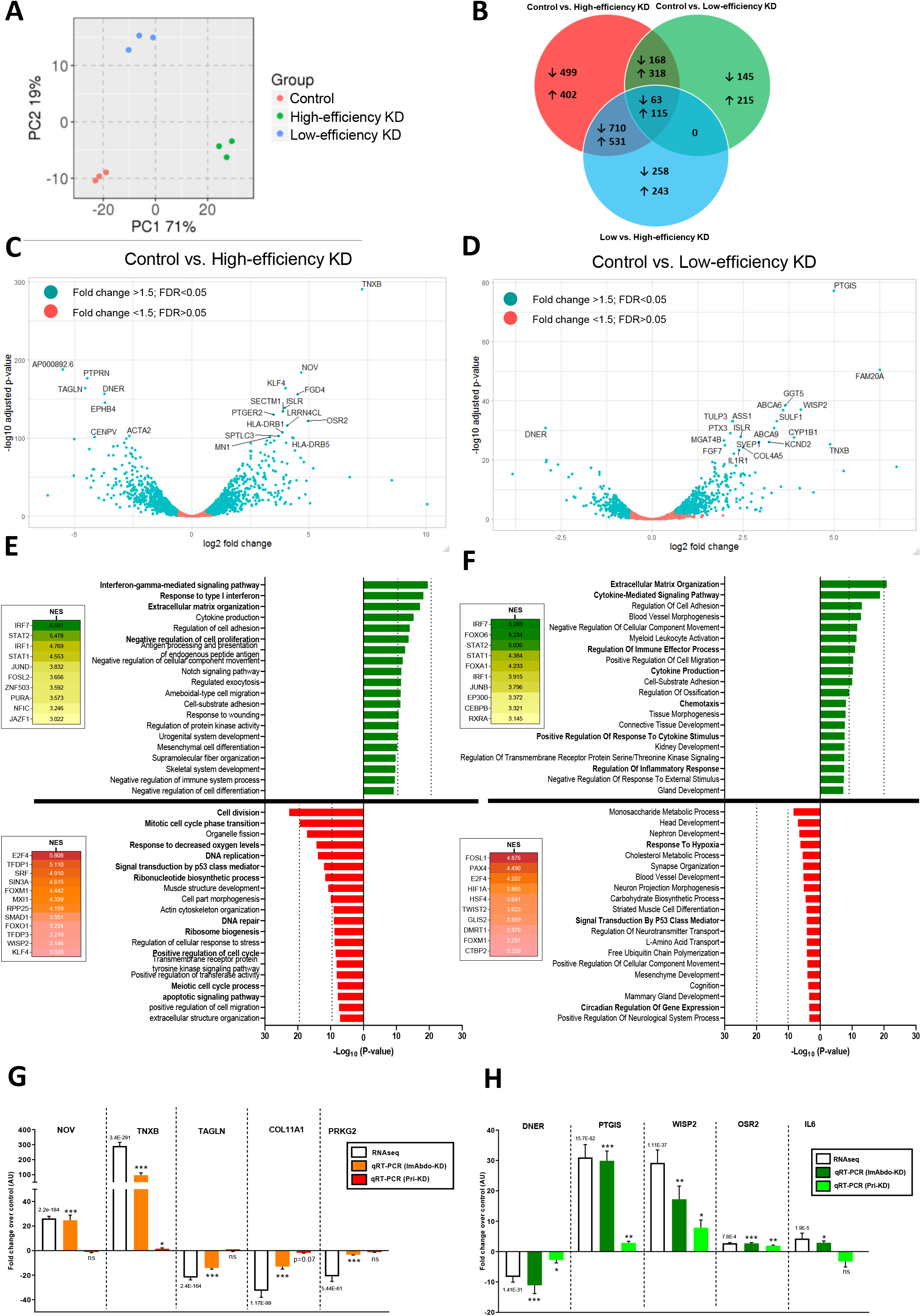
Global transcriptional profiling reveals that TCF7L2 regulates multiple aspects of AP biology. (A) Principal component analysis (PCA) of the global transcriptomic profile of control (scrambled), moderate (sh897) and high (sh843) TCF7L2 KD DFAT abdominal APs. (B) Venn diagram showing the overlap between differentially regulated genes from paired comparisons (FDR<0.05; absolute fold change >1.5). (C and D) Volcano plots showing the genes differentially regulated between control and (C) sh897; or (D) sh843, TCF7L2 KD DFAT abdominal APs. Highlighted are the top 20 differentially regulated genes. (E and F) Pathway enrichment analyses of genes downregulated (red) and upregulated (green) in (E) sh897 and (F) sh843 DFAT abdominal APs, *vs*. controls. Shown also are transcription factor binding-site motif analysis (inset) of downregulated (red) and upregulated (green) genes. (G and H) qRT-PCR validation of differentially regulated genes. qRT-PCR data were normalized to 18S rRNA levels and are from 3 independent experiments. Histograms are means ± SEM. FDR is annotated for RNA-Seq fold-change measurements. ***p<0.001, **p<0.01, *p<0.05 for qRT-PCR measurements.

### The T2D-risk allele at rs7903146 reduces AP TCF7L2 expression

Genome wide association study (GWAS) meta-analyses have revealed that non-coding genetic variation at *TCF7L2* is the strongest genetic determinant of T2D risk in humans (16,17). Previous efforts to characterize the mechanism of action at *TCF7L2* revealed that the fine-mapped T2D-risk allele at rs7903146 overlaps enhancer histone marks in both islets and APs (30). Whilst this variant has been shown to increase *TCF7L2* expression in islets (31), evidence demonstrating that it is associated with a cis-expression quantitative trait locus (eQTL) in APs has been missing. By examining age-, BMI- and sex-adjusted *TCF7L2* mRNA abundance data in isolated mADs and *ex vivo* cultured abdominal and gluteal APs from up to 21 homozygous carriers of the T2D-risk variant (T) and 59 subjects homozygous for the wild-type allele (C) at rs7903146 we found that the T allele reduces *TCF7L2* expression in abdominal APs (Fig. 6A, B). A similar trend was detected in gluteal APs. We substantiated this finding in a subset of samples by demonstrating lower TCF7L2 protein levels in abdominal APs derived from T2D-risk variant carriers *versus* ancestral allele carriers (Fig. 6C). In cell-based luciferase assays a 150 base pair nucleotide genomic sequence centred around rs7903146 exhibited allele-specific enhancer properties in abdominal APs and HEK293 cells with the T allele abolishing enhancer activity (Fig. 6E, F). Histological assessment of abdominal AT biopsies from 19 age- and BMI-matched pairs of males homozygous either for the T2D-risk allele or the protective allele at rs7903146 did not reveal any difference in median adipocyte size between the two genotypes (Fig. 6G). However, interrogation of plasma biochemistry data from 600 age- and BMI-matched pairs of subjects from the Oxford Biobank showed that T2D-risk variant carriers had reduced AT insulin resistance (Adipo-IR) (Fig. 6H) which was more pronounced in obese individuals (Fig. 6I). We conclude that in addition to its established role in regulating pancreatic insulin secretion, genetic variation at rs7903146 may also influence T2D risk through effects on AP biology.

**Fig. 6.**
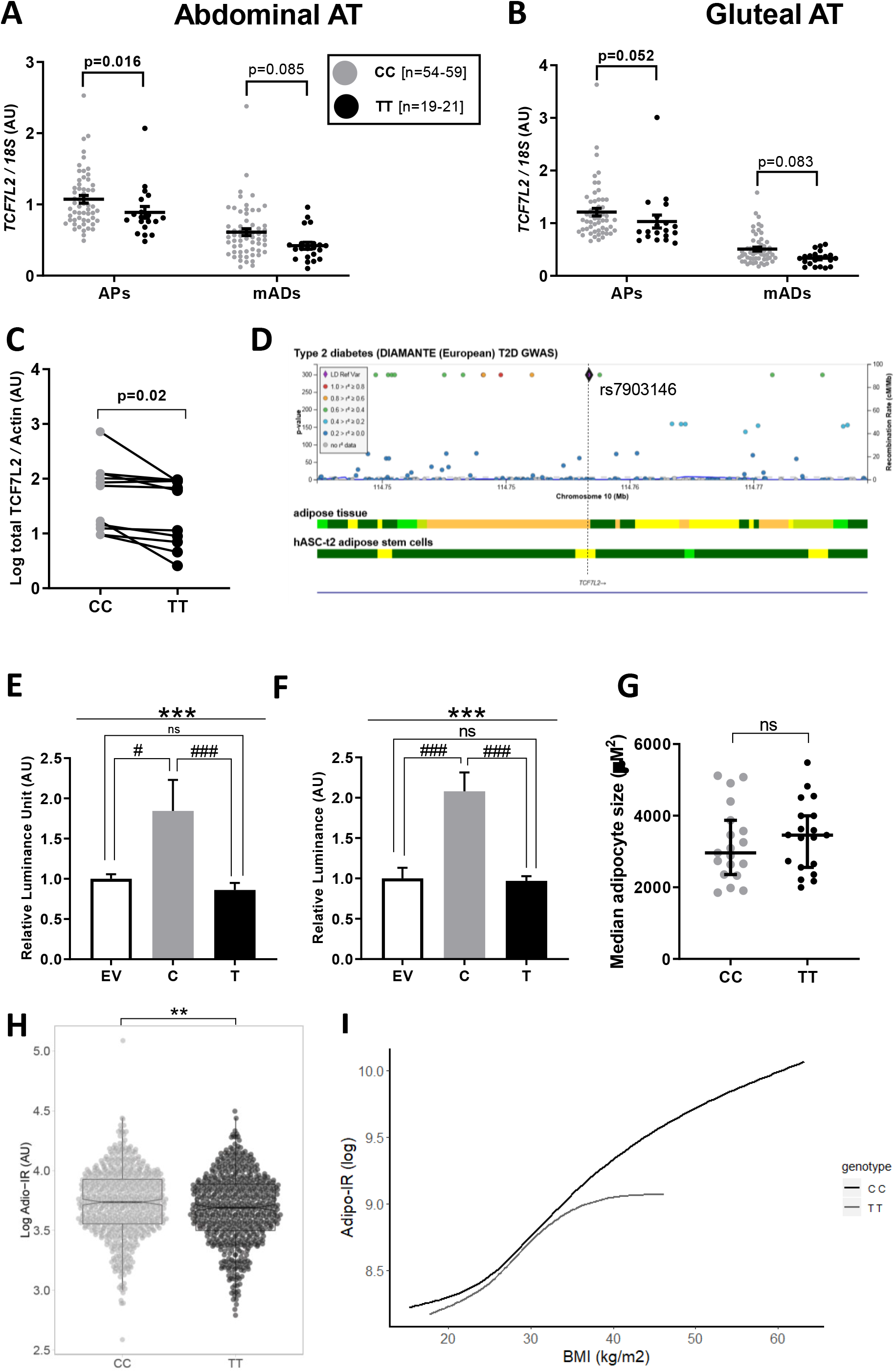
The type 2 diabetes risk allele at rs7903146 reduces TCF7L2 expression in human abdominal APs. (A and B) *TCF7L2* expression in paired cultured APs and mature adipocytes (mADs) from subcutaneous (A) abdominal and (B) gluteal AT biopsies from homozygous carriers of the T2D allele T (n=19-21 [4F]) *vs*. carriers of the non-risk allele C (n=54-59 [29F]). TT carriers, Age - 42.8±6.4 years, BMI - 26.2±6.0 kg/m^2^. CC carriers, Age - 45.8±9.5 years, BMI - 27.3±3.8 kg/m^2^. p-value adjusted for age, BMI and sex. (C) TCF7L2 protein levels in cultured abdominal APs from age-, BMI- and sex-matched homozygous carriers of the T2D risk allele T *vs*. carriers of the non-risk allele C (n=11/group). Actin was used as western blot loading control. (D) Chromatin state map showing that rs7903146 overlaps a weak enhancer in APs (image reproduced from type2diabetesgenetics.org). Yellow – weak enhancer, orange – active enhancer, green – transcription (E and F) Luciferase reporter assay in (E) DFAT abdominal APs and (F) HEK293 cells transfected with empty vector (EV) or the luciferase reporter vector containing 151 bp genomic sequence with the T2D risk allele T or the non-risk allele C (n=12-16 replicates/group). ***p<0.001; ###p<0.001, #p<0.05 (adjusted for multiple comparisons). (G) Median adipocyte area (μm^2^) calculated from the histological sections of abdominal AT from 19 pairs of age- and BMI-matched males grouped by rs7903146 genotype. CC carriers, Age - 44.2±6.3 years, BMI - 25.4±2.8 kg/m^2^. TT carriers, Age - 44.1±7.3 years, BMI - 25.1±2.7 kg/m^2^. Error bars are median with 95% CI. (H) Comparison of adipose tissue insulin resistance (Adipo-IR) between age-, BMI- and sex-matched homozygous carriers of the T2D risk allele (T) and the non-risk allele (C) (n= 600 pairs). **p<0.01. Data are medians with boxplot showing IQR and whiskers are range of values within 1.5*IQR. (I) Smoothened splines showing the relationship between adipo-IR and BMI for homozygous carriers of the T2D risk allele (T) and carriers of the non-risk allele (C) at rs7903146. See Table S1 for anthropometric and plasma biochemistry data for (H and I).

## DISCUSSION

Our study highlights a complex role for TCF7L2 in AP biology and AT function. We show that *TCF7L2* expression is similar across different fat depots and diminishes in abdominal AT in obesity. The lower AT *TCF7L2* mRNA abundance in obese subjects could be mediated via increased methylation at *TCF7L2* (32) and may contribute to the AT dysfunction that is associated with greater adiposity. Our results also revealed that *TCF7L2* was highly expressed in all adipose cell lineages examined, namely APs, mature adipocytes and endothelial cells with the former exhibiting the highest transcript levels. These data suggest that TCF7L2 has pleiotropic roles in AT ranging from the regulation of AP biology and adipogenesis, to the modulation of mature adipocyte function and angiogenesis. Finally, we showed that increased adiposity was paradoxically associated with higher *TCF7L2* expression in isolated abdominal APs despite reduced *TCF7L2* mRNA abundance in whole AT. It is likely that the reduction in *TCF7L2* mRNA levels in the abdominal depot in obesity is driven by lower expression in adipocytes given that they are the principal cellular component of AT and as hinted by the correlation data in Fig. 1F. Thus, obesity seems to lead to directionally opposite changes in *TCF7L2* expression in different AT cellular fractions.

To elucidate the role of TCF7L2 in AP biology, we undertook functional studies in immortalised and primary APs. These revealed that TCF7L2 protein production may be regulated both transcriptionally and at the level of protein translation because TCF7L2-KD was associated with more pronounced changes in protein than mRNA levels. Thus in addition to regulating its own transcription (33) TCF7L2 may regulate its own translation. The loss-of-function studies also showed that TCF7L2-KD impairs AP proliferation. Additionally, they demonstrated that TCF7L2 dose-dependently regulates adipogenesis. We speculate that the primary role of TCF7L2 in APs is to restrain adipogenesis since only complete TCF7L2-KD led to impaired adipocyte differentiation whilst downregulation of its expression within a physiological range, in both DFAT and primary abdominal APs, led to enhanced adipogenesis.

To decipher the signalling pathways mediating the actions of TCF7L2 in AP biology we examined canonical WNT signalling in TCF7L2-KD cells. These experiments revealed that in abdominal APs TCF7L2 may function to inhibit WNT pathway activity. Whilst *prima facie* paradoxical, this result is in keeping with previous findings demonstrating that TCF7L2-KD in 3T3L1 cells was associated with a robust increase in *Axin2* expression (11). Tang *et al* (34) also showed that TCF7L2 displayed a cell-type specific activity to both enhance and inhibit WNT signalling and localized the transcriptional repressive ability of TCF7L2 to its C-terminal tail, which is present in some, so-called E isoforms, but not all *TCF7L2* transcript species. However, expression of E-type *TCF7L2* splice variants is low in AT (35,36). Additionally, none of these studies examined the expression of other TCF/LEF family members in TCF7L2-KD cells (see below). WNT/β-catenin signalling is known to inhibit adipogenesis. Accordingly, high-level KD of TCF7L2 was associated with impaired adipogenesis concomitant with canonical WNT signalling activation. However, downregulation of TCF7L2 expression within a more physiological range (using sh897) led to increased adipocyte differentiation despite mild WNT/β-catenin pathway activation. These data suggest that WNT signalling may have dose-dependent effects on adipogenesis and cell fate determination (21,37). Additionally/alternatively TCF7L2 may engage other signalling pathways to influence AP biology.

To further elucidate the genes, pathways and biological processes regulated by TCF7L2 in APs we undertook genome-wide transcriptional profiling of TCF7L2-KD cells. Consistent with our *in vitro* studies these experiments showed that KD of TCF7L2 activates transcriptional programmes which inhibit the proliferation and modulate the adipogenic capacity of APs. Furthermore, they revealed that TCF7L2 can regulate many aspects of AP biology including ECM secretion, immune and inflammatory signalling, and apoptosis and/or senescence. TCF7L2-KD in APs also led to suppression of HIF1A signalling which promotes fibrosis and AT dysfunction (3). These findings expand the potential functional repertoire of TCF7L2 in AT although, they require experimental confirmation. Contrary to our expectations, the RNA-Seq experiments also revealed that TCF7L2 promotes canonical WNT signalling in APs. How can we reconcile this finding with our *in vitro* studies? In TCF7L2-KD cells expression of *TCF7* was upregulated. We speculate that TCF7 is a more potent activator of WNT signalling in APs given that functional redundancy is common amongst TCF/LEF family members (8) and based on the increased TOPflash promoter activity and *AXIN2* expression in TCF7L2-KD cells. Nonetheless, increased TCF7 levels cannot compensate for the absence of TCF7L2 at many WNT target gene promoters. Consequently, AP TCF7L2-KD leads to downregulation of several classic WNT target genes and pathways. Transcription factor-binding site analysis was in keeping with the results of the gene set enrichment analysis. Interestingly we saw no enrichment for TCF7L2 binding in the promoters of genes suppressed in TCF7L2-KD cells. However, genome-wide chromatin occupancy data have shown that TCF7L2 regulates gene expression primarily by binding to intergenic regions (29,33). Lastly, the RNA-Seq data provided additional evidence that partial TCF7L2-KD in DFAT cells may be more physiologically relevant as we were better able to replicate changes in the transcriptome of partial *versus* complete TCF7L2-KD DFAT cells in moderate- and high-efficiency TCF7L2-KD primary cells respectively.

GWAS meta-analyses have identified eight independent signals at *TCF7L2* which are associated with T2D susceptibility (17). At least three of these overlap regions of active chromatin in AT (17). Of these signals, only rs7903146 has received attention to date. It has been demonstrated through analysis of GWAS metadata of T2D-related traits (38,39) and human physiological studies (40) that the risk allele at this SNV increases T2D susceptibility *via* impaired insulin secretion. Additionally, this variant was shown to increase *TCF7L2* expression in pancreatic islets (31,41). These data have conclusively established that rs7903146 increases T2D risk primarily through islet dysfunction probably driven by increased *TCF7L2* expression in pancreatic beta-cells. Nevertheless, this SNV may also influence T2D predisposition *via* actions in AT as it was shown to overlap active enhancer histone marks in APs (30) and to be associated with fat distribution (18). Using a “soft clustering” method to group variant-trait associations ascertained from GWAS for 94 independent T2D signals and 47 diabetes-related traits another study also provided some evidence that “lipodystrophy-like” insulin resistance may contribute to T2D predisposition at rs7903146 (39). The same SNV was further shown to overlap a DNA-methylated genomic region exhibiting increased methylation in AT from obese *versus* lean subjects which was reversed post gastric bypass surgery (32). Extending these findings, we now demonstrate that rs7903146 is an eQTL for *TCF7L2* in abdominal APs with the T allele *reducing TCF7L2* expression. Whilst the eQTL signal at rs7903146 was only nominally significant our gene expression data were derived from *ex vivo* expanded APs. This would have added noise to the eQTL dataset in addition to the noise introduced by inter-individual variability in *TCF7L2* AP expression. We also corroborated our data by demonstrating that the T2D-risk variant at this SNV was associated with reduced AP TCF7L2 protein levels and displayed allele specific enhancer activity in luciferase assays which was directional consistent with the eQTL data. Interestingly rs7903146 confers higher T2D susceptibility in lean *versus* obese subjects (17). Our results offer a possible explanation for this paradox. In our study the T2D-risk variant at this SNV was associated with enhanced age- and BMI-adjusted AT insulin sensitivity which was more pronounced in obese subjects. Whilst we detected no difference in adipocyte size based on rs7903146 genotype, our study sample (n=19) comprised mostly of lean (n=7) and overweight subjects (n=11) due to limited numbers of obese homozygous T2D-risk allele carriers in the Oxford Biobank (see Fig. 6I). Based on the dose-dependent effects of TCF7L2 on adipogenesis demonstrated herein, it will be interesting to determine whether the T2D-predisposing allele at rs7903146 is associated with differential effects on adipocyte size in lean *versus* obese subjects; with smaller fat cells specifically seen in obese individuals which have higher baseline AP *TCF7L2* expression.

In summary, our work highlights that TCF7L2 plays an important but complex role in AP biology. We also demonstrate that rs7903146 has *cis* regulatory effects outside the pancreas and reduces *TCF7L2* expression in abdominal APs. Thus, in addition to islet dysfunction, altered AP function consequent to changes in *TCF7L2* expression might also influence the T2D predisposition conferred by this SNV. Future studies should define the role of TCF7L2 in human adipocytes and characterize the impact of T2D-associated regulatory variants at *TCF7L2*, beyond rs7903146, which overlap open chromatin in adipose cells on adipose *TCF7L2* expression and AT function.

## Supporting information

Online supplemental material_Verma et al., 2019

## ACKNOWLEDGEMENTS

We thank the volunteers from the Oxford BioBank (www.oxfordbiobank.org.uk) for their participation in this recall study. We thank the nurses at the Clinical Research Unit, in particular, Mrs. Jane Cheeseman, for their help in recruitment and sample collection from Oxford BioBank volunteers. The OBB and Oxford BioResource are funded by the NIHR Oxford Biomedical Research Centre (BRC). The views expressed are those of the author(s) and not necessarily those of the NIHR or the Department of Health and Social care. This research is supported by the British Heart Foundation through an Intermediate Clinical Research Fellowship to CC (FS/16/45/32359) and a programme grant (RG/17/1/32663) to FK. MV and ADVD are supported by a Novo Nordisk Postdoctoral Fellowship run in partnership with the University of Oxford. Funding support was also received from the National Institute for Health Research, Oxford Biomedical Research Centre (BRC). We thank the Oxford Genomics Centre at the Wellcome Centre for Human Genetics (funded by Wellcome Trust grant reference 203141/Z/16/Z) for the generation of the Sequencing data. The Genotype-Tissue Expression (GTEx) Project was supported by the Common Fund of the Office of the Director of the National Institutes of Health, and by NCI, NHGRI, NHLBI, NIDA, NIMH, and NINDS. The data used for the analyses described in this manuscript were obtained from the GTEx Portal on 10/20/19.

## CONTRIBUTION STATEMENT

Conceptualization, C.C.; Methodology, M.V., N.Y.L., S.K.V.; Investigation, M.V., N.Y.L., S.K.V., A.D.V.D., M.T., M.J.N., C.C., Writing – Original Draft, M.V., C.C.; Writing – Review & Editing, All authors; Funding Acquisition, C.C., F.K., Resources, C.C., F.K.; Supervision, C.C., F.K. C.C is the guarantor of this work and, as such, had full access to all the data in the study and takes responsibility for the integrity of the data and the accuracy of the data analysis.

## PRIOR PRESENTATIONS

Parts of the study were presented as an oral presentation at the 54^th^ Annual Meeting of European Association for the Study of Diabetes, Berlin, Germany, 1-5^th^ October 2018 and at the 44^th^ Adipose Tissue Discussion Group meeting, Edinburgh, UK, 6-7^th^ December 2018.

