## Supplementary material for "TCF7L2 plays a complex role in human adipose progenitor biology which may contribute to genetic susceptibility to type 2 diabetes": Online supplemental material_Verma et al., 2019

#### SUPPLEMENTARY FIGURE LEGENDS

**Fig. S1 Correlation between *TCF7L2* expression in human gluteal adipose cell fractions and donor body mass index (BMI).** (A) Correlation between *TCF7L2* expression in cultured APs from subcutaneous gluteal AT and donor BMI (n=107 [59F] / group; Age - 45.0±8.4 years, BMI - 27.2±4.5 kg/$m^2$). (B) Correlation between *TCF7L2* expression in mADs from subcutaneous gluteal AT and donor BMI. (n=110 [59F] / group; Age - 45.1±8.5 years, BMI - 27.0±4.1 kg/$m^2$). qRT-PCR data were normalized to 18S rRNA levels.

**Fig. S2 DFAT APs maintain their depot-specific gene expression pattern *in vitro*.** (A) *HOXA6* and *HOXC11* mRNA levels in DFAT abdominal and gluteal APs. (B) *HOXA6*, *HOXA5*, *SHOX2* and *HOTAIR* mRNA levels in DFAT abdominal and gluteal APs at final day 15 of adipogenic differentiation. qRT-PCR data were normalized to 18S rRNA levels. Histograms are means ± SEM. ***p<0.001, **p<0.01.

**Fig. S3 Low dose TCF7L2 overexpression does not alter adipogenesis in DFAT abdominal APs.** (A) Representative micrographs of DFAT abdominal APs stably transduced with the empty vector (EV) or TCF7L2 overexpression vector (TCF7L2), and cultured in the presence of vehicle (Veh) or 0.01 µg/ml DOX, at day 14 of the adipogenic differentiation. The histogram shows the relative lipid accumulation, assessed by AdipoRed lipid stain (n=16 wells/group). (B - E) Relative mRNA levels of adipogenic genes *CEBPA*, *PPARG2*, *FABP4* and *ADIPOQ* at day 14 of adipogenic differentiation. qRT-PCR data were normalized to 18S rRNA levels. Histograms are means ± SEM and expressed relative to vehicle treatment (arbitrarily set to 1) for EV or TCF7L2 overexpressing DFAT abdominal AP line, respectively. Data obtained from at least 3 independent experiments. ***p<0.001, **p<0.01.

**Fig. S4 Confirmation of *TCF7L2* overexpression in *in vitro* differentiated DFAT abdominal cells.** DFAT abdominal APs stably transduced with either the empty vector (EV) or TCF7L2 overexpression vector (TCF7L2) were cultured in the presence of vehicle (Veh) or (0.01 µg/ml or 0.05 µg/ml) DOX to induce TCF7L2 overexpression. *TCF7L2* expression was determined by qRT-

PCR in *in vitro* differentiated cells at (A) day 7, or (B) day 14 of adipogenic differentiation. qRT-PCR data were normalized to 18S rRNA levels. Histograms are means ± SEM and expressed relative to vehicle-treated (arbitrarily set to 1) EV or TCF7L2 overexpressing DFAT abdominal APs, respectively. Data obtained from at least 3 independent experiments. ***p<0.001, **p<0.01, *p<0.05; $^{\#\#\#}$p<0.001, $^{\#}$p<0.05 (adjusted for multiple comparisons).

**Fig. S5 High dose TCF7L2 overexpression accelerates adipogenesis in DFAT abdominal APs.** DFAT abdominal APs stably transduced with either the empty vector (EV) or TCF7L2 overexpression vector (TCF7L2) were cultured in the presence of vehicle (Veh) or 0.05 µg/ml DOX. (A) Representative micrographs at day 7 of adipogenic differentiation. The histogram shows the relative lipid accumulation, assessed by AdipoRed lipid stain (n=16 wells/group). (B - E) Relative mRNA levels of adipogenic genes *CEBPA*, *PPARG2*, *FABP4* and *ADIPOQ* at day 7 of adipogenic differentiation. (F) Representative micrographs at day 10 of adipogenic differentiation. The histogram shows the relative lipid accumulation, assessed by AdipoRed lipid stain (n=16 wells/group). qRT-PCR data were normalized to 18S rRNA levels. Histograms are means ± SEM and expressed relative to vehicle-treated (arbitrarily set to 1) EV or TCF7L2 overexpressing DFAT abdominal APs, respectively. Data obtained from at least 3 independent experiments. ***p<0.001, **p<0.01.

**Fig. S6 *TCF7L1* levels are unaltered with knockdown or overexpression of TCF7L2 in DFAT abdominal APs.** (A) *TCF7L1* mRNA levels in control (scrambled), moderate (sh897) and high (sh843) TCF7L2 KD DFAT abdominal APs. (B) *TCF7L1* mRNA levels in DFAT abdominal APs transduced with either the empty vector (EV) or TCF7L2 overexpression vector (TCF7L2), cultured in the presence of either vehicle (Veh) or DOX (0.01 µg/ml or 0.05 µg/ml). Data expressed relative to vehicle-treated (arbitrarily set to 1) EV or TCF7L2 overexpressing DFAT abdominal AP line, respectively. qRT-PCR data were normalized to 18S rRNA levels. Histograms are means ± SEM. Data obtained from at least 3 independent experiments. *p<0.05.

**Fig. S7 Expression profile of direct WNT target genes following TCF7L2 KD in DFAT abdominal APs.** $Log_2$ fold change in mRNA levels of differentially regulated (FDR<0.05) direct

WNT target genes in control (scrambled) *vs.* moderate (sh897) or control (scrambled) *vs.* high (sh843) TCF7L2 KD DFAT abdominal APs as measured by RNA-Seq analysis.

**Supplementary Table. 1 Anthropometry and plasma biochemistry measurements in homozygous carriers of type 2 diabetes risk-allele (T) or the non-risk allele (C).** (A) Age and BMI measurements in age-, BMI-, and sex-matched homozygous carriers of the type 2 diabetes risk-allele (T) or the non-risk allele (C). (B) Anthropometric and plasma biochemistry measurements in homozygous carriers of the type 2 diabetes risk-allele (T) and the non-risk allele (C) in the Oxford Biobank.

#### SUPPLEMENTAL METHODS:

**Endothelial (CD31$^{+}$) cell isolation from adipose tissue:** Stromal-vascular cells from human adipose tissue biopsies were obtained as described previously (1), resuspended in Endothelial Basal Medium-2 with Growth Supplements (EGM-2 Endothelial Medium BulletKit, Lonza) and seeded into 1% gelatin-coated flasks. Cells were expanded until a confluent T75 flask was obtained. Upon trypsinization, cells were resuspended in 90uL PBS + 2mM EDTA + 0.5% BSA (PEB) buffer. Enrichment of CD31$^{+}$ cells was done by magnetic-activated cell sorting (MACS) using MiniMACS Separation columns and human CD31 MicroBead Kit (all from Miltenyi Biotec) following manufacturers' protocol for isolation of endothelial cells from mouse skeletal muscle. Both the CD31$^{-}$ (AP) and CD31$^{+}$ (endothelial cell) fractions were collected and seeded into 6-well and 1% gelatin-coated 24-well plates respectively. Media was changed the following day to remove any remaining PEB buffer. Cells were cultured until confluent, after 3 to 4 days, and harvested for RNA extraction.

**Generation of DFAT APs:** DFAT cells were generated by selection and de-differentiation of lipid-laden, *in vitro* differentiated immortalised APs (2) with modifications. Briefly, immortalised APs were differentiated, and at the intermediate time point (day 7) of adipogenic differentiation, (50ng/ml) recombinant human BMP2, and 180μM fatty acid mix (oleate 75μM, palmitate 65μM, linoleate 40μM dissolved in 5% BSA) were added. Well differentiated, detached, and lipid-laden cells in the culture media were transferred to 9cm$^2$ slide flasks filled with standard AP growth media for ceiling culture, and allowed to adhere and de-differentiate for 3-5 days. The side flasks were then inverted, and fresh standard AP growth media was added to allow for de-differentiation. These daughter DFAT APs showed higher adipogenic capacity than immortalised and primary APs.

**RNA isolation and real time-PCR:** Total RNA from APs and *in vitro* differentiated APs were extracted by using RNAeasy mini kit (Qiagen) and from AT biopsies using TRIzol® reagent (Invitrogen). 0.5 and 1.0μg (for APs or *in vitro* differentiated APs) or 0.2μg (mADs) of purified RNA were reverse transcribed into cDNA using the High Capacity cDNA Reverse Transcription kit (Applied Biosystems). TaqMan gene expression assays were used for qRT-PCR as described (3) (see the table below for TaqMan assay IDs). mRNA expression levels were normalised to geometric mean of either *PPIA*, *PGK1*, *PSMB6* and *IPO8*, or *PPIA* and *PGK1* for AT samples or to 18S rRNA levels for cultured cells.

| **Gene** | **TaqMan Assay** |
|---|---|
| *TCF7L2* | Hs01009044_m1 |
| *18S* | Hs99999901_s1 |
| *PPIA* | Hs99999904_m1 |
| *PGK1* | Hs99999906_m1 |
| *PSMB6* | Hs00382586_m1 |
| *IPO8* | Hs00183533_m1 |
| *AXIN2* | Hs00610344_m1 |
| *CEBPA* | Hs00269972_s1 |
| *PPARG2* | Hs01115510_m1 |
| *FABP4* | Hs00609791_m1 |
| *ADIPOQ* | Hs00605917_m1 |
| *TCF7* | Hs01556515_m1 |
| *TCF7L1* | Hs01064103_m1 |
| *LEF1* | Hs01547250_m1 |

| | |
|---|---|
| *NOV* | Hs00159631_m1 |
| *TNXB* | Hs00372889_g1 |
| *TAGLN* | Hs01038777_g1 |
| *COL11A1* | Hs01097664_m1 |
| *PRKG2* | Hs00211903_m1 |
| *DNER* | Hs01039911_m1 |
| *PTGIS* | Hs00919949_m1 |
| *WISP2* | Hs01031984_m1 |
| *OSR2* | Hs01085594_m1 |
| *IL6* | Hs00174131_m1 |

**Western blots:** Preparation of whole cell lysates, quantification of protein, western blotting and densitometry for protein levels were done as described (1). Details of the antibodies are listed in below.

| Antibody | Supplier |
|---|---|
| anti-TCF7L2 | Clone C48H11 #2569, Cell Signaling Technology |
| anti-Active β-CATENIN | Clone 8E7, 05-665, Merck Millipore |
| anti-α-TUBULIN | ab15246, Abcam |
| anti-ACTIN | sc-1616, Santa Cruz Biotechnology |

**RNA sequencing:** RNA sequencing was performed on scrambled controls, sh897 and sh843 TCF7L2 KD DFAT abdominal APs from 3 independent experiments. Extraction of total RNA and on-column DNAseI digestion was performed using RNeasy mini kit (Qiagen). RNA quantification and QC was done on NanoDrop ND-1000 (Labtech) and Agilent 2100 Bioanalyzer (Agilent) and the 2200 or 4200 Tape Station (Agilent, RNA ScreenTape), respectively. RNA integrity number (RIN) estimates were ≥8 for all samples. Library preparation and cDNA sequencing was done at the Oxford Genomics Centre (Wellcome Trust Centre for Human Genetics, Oxford, UK). Briefly, strand specific library preparation was undertaken following polyadenylated transcript enrichment using NEB Next Ultra II mRNA kit (NEB). Unique dual indexing primers were utilised for library amplification on Tetrad (Bio-Rad) (4). Library quantification and profile analysis was done on Qubit (Thermo Fischer) and the 2200 or 4200 Tape Station (Agilent, RNA ScreenTape), respectively. Individual libraries were normalized, pooled and diluted to ~10nM for storage. The 10nM library was denatured, diluted and paired end sequenced on HiSeq4000 75bp platform (Illumina, HiSeq 3000/4000 PE Cluster Kit and 150 cycle SBS Kit) which generated >30 million reads per sample. Bioinformatic analysis of raw sequencing data was undertaken by Cambridge Genomic Services (Cambridge, UK). Briefly, read QC, trimming and mapping was done by FastQC (5), TrimGalore (6) and STAR (7) (using Ensembl Homo_sapiens.GRCh38 (release 83) as reference genome), respectively. Read counting was undertaken using HTSeq (8). Principal Component Analysis (PCA) plots were based on normalised and rlog transformed counts obtained with DESeq2 (9). Differential gene expression analysis was done using the counted reads and the R package edgeR (10) for the 3 pairwise comparisons (Control (scrambled) *vs.* Low-efficiency KD, Control (scrambled) *vs.* High-efficiency KD and Low-efficiency KD *vs.* High-efficiency KD). Genes with very low expression were excluded based on the principle that a gene has to be expressed at the equivalent ≥ 5 reads (CPM) in at least 3 samples when using all 6 samples in the pairwise comparisons. Adjustment for multiple testing was undertaken *via* the FDR (Benjamini-Hochberg) approach used by edgeR. Pathway enrichment and transcription factor binding-site motif analysis was done in differentially expressed genes (FDR <0.05, absolute fold change >1.5) using Metascape (11) and iRegulon (12), respectively.

throughput library preparation using the transposome-based nextera system. BMC Biotechnol. 2013;13.

**Fig. S1**

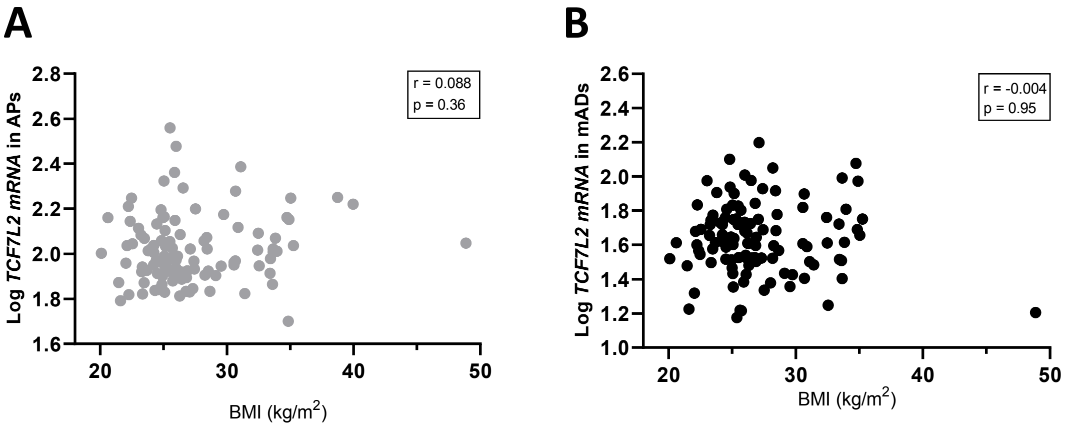
A
B
Log TCF7L2 mRNA in APs
r = 0.088
p = 0.36
2.8
2.6
2.4
2.2
2.0
1.8
1.6
20
30
40
50
BMI (kg/m2)
Log TCF7L2 mRNA in mADs
r = -0.004
p = 0.95
2.6
2.4
2.2
2.0
1.8
1.6
1.4
1.2
1.0
20
30
40
50
BMI (kg/m2)

**Fig. S2**

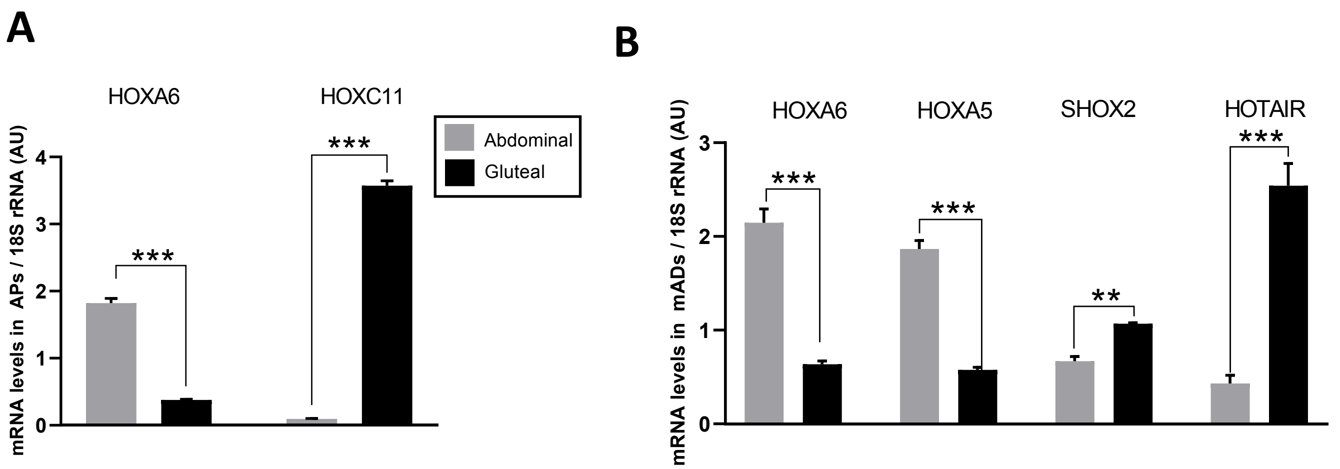
A
HOXA6
HOXC11
mRNA levels in APs / 18S rRNA (AU)
0
1
2
3
4
***
***
Abdominal
Gluteal
B
HOXA6
HOXA5
SHOX2
HOTAIR
mRNA levels in mADs / 18S rRNA (AU)
0
1
2
3
***
***
**
***

Fig. S3

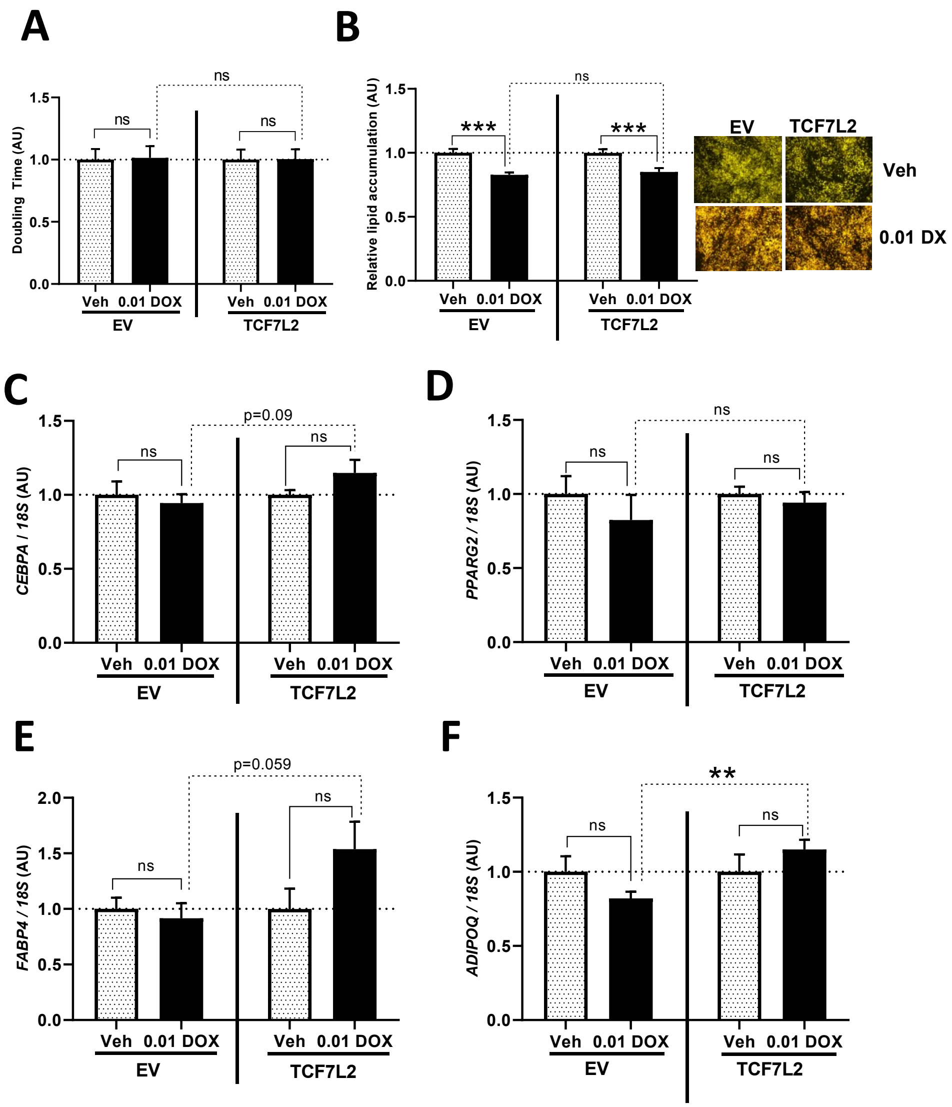
A
B
C
D
E
F
Doubling Time (AU)
Relative lipid accumulation (AU)
CEBPA / 18S (AU)
PPARG2 / 18S (AU)
FABP4 / 18S (AU)
ADIPOQ / 18S (AU)
ns
***
p=0.09
p=0.059
**
Veh 0.01 DOX
EV
TCF7L2
Veh
0.01 DX

**Fig. S4**

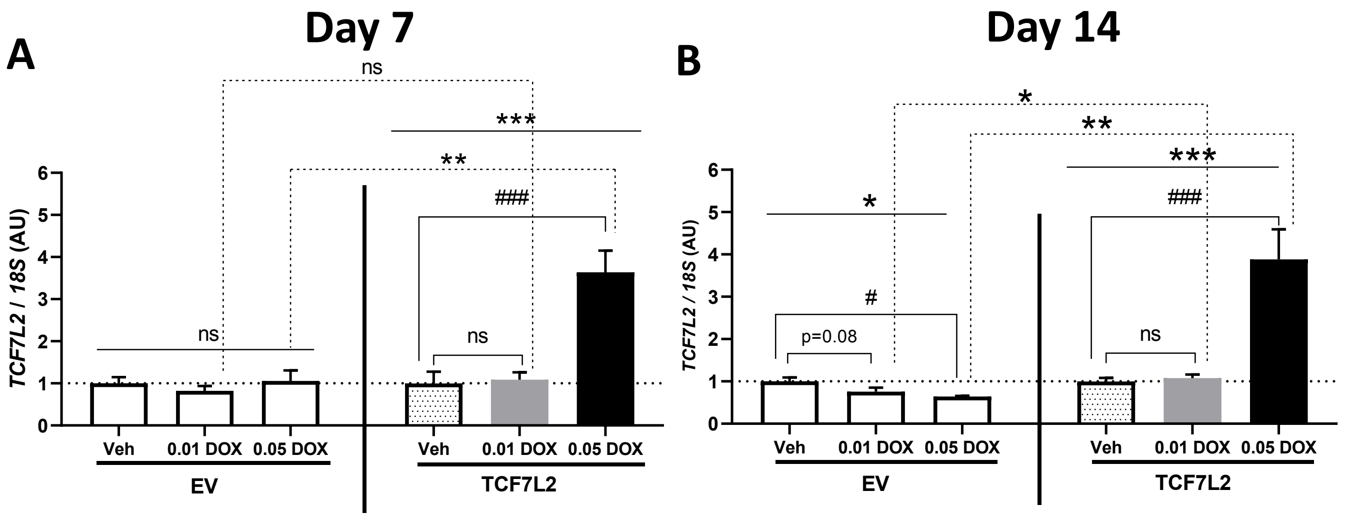
A
Day 7
B
Day 14
TCF7L2 / 18S (AU)
ns
***
**
###
Veh
0.01 DOX
0.05 DOX
EV
TCF7L2
*
#
p=0.08

### Fig. S5

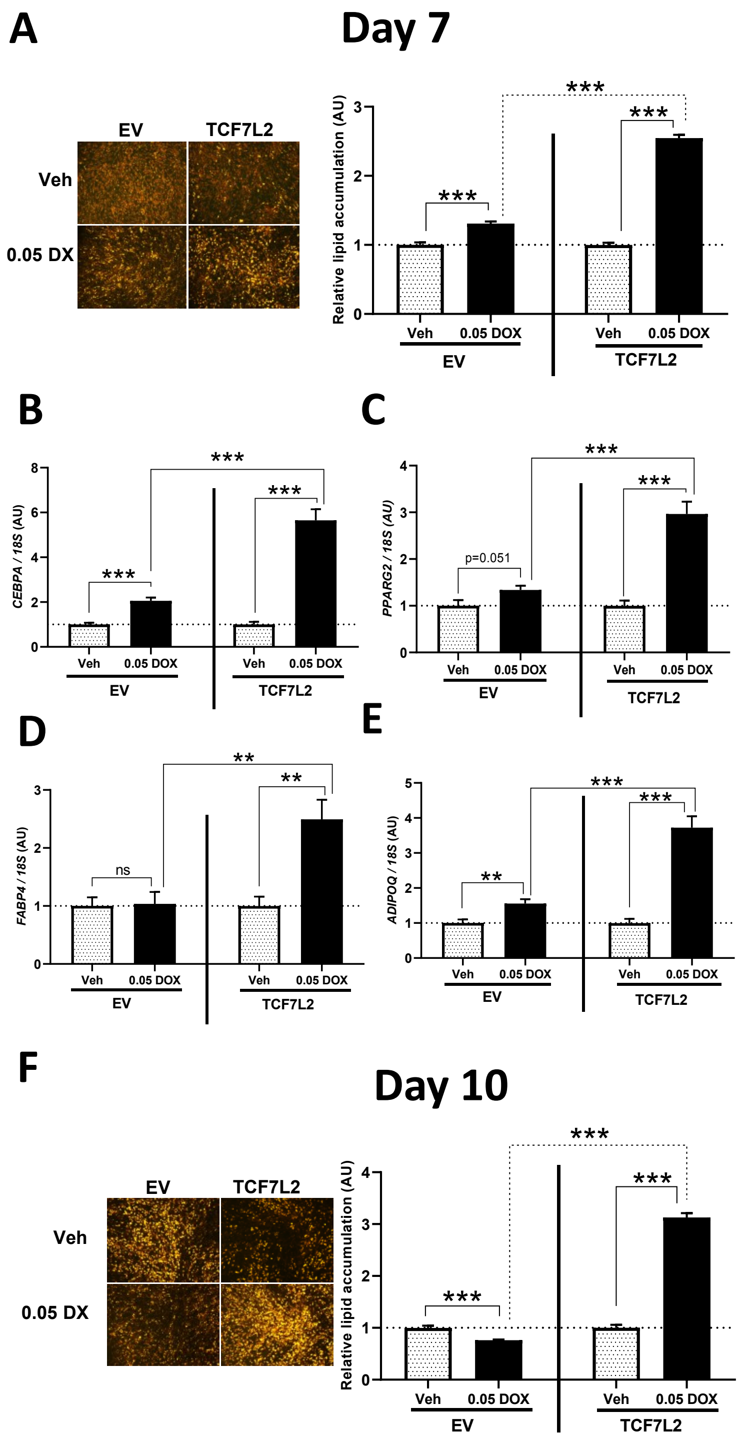

A
Day 7
EV
TCF7L2
Veh
0.05 DX
Relative lipid accumulation (AU)
***
***
***
Veh
0.05 DOX
Veh
0.05 DOX
EV
TCF7L2
B
CEBPA / 18S (AU)
***
***
***
Veh
0.05 DOX
Veh
0.05 DOX
EV
TCF7L2
C
PPARG2 / 18S (AU)
***
***
p=0.051
Veh
0.05 DOX
Veh
0.05 DOX
EV
TCF7L2
D
FABP4 / 18S (AU)
**
**
ns
Veh
0.05 DOX
Veh
0.05 DOX
EV
TCF7L2
E
ADIPOQ / 18S (AU)
***
***
**
Veh
0.05 DOX
Veh
0.05 DOX
EV
TCF7L2
F
Day 10
EV
TCF7L2
Veh
0.05 DX
Relative lipid accumulation (AU)
***
***
***
Veh
0.05 DOX
Veh
0.05 DOX
EV
TCF7L2

**Fig. S6**

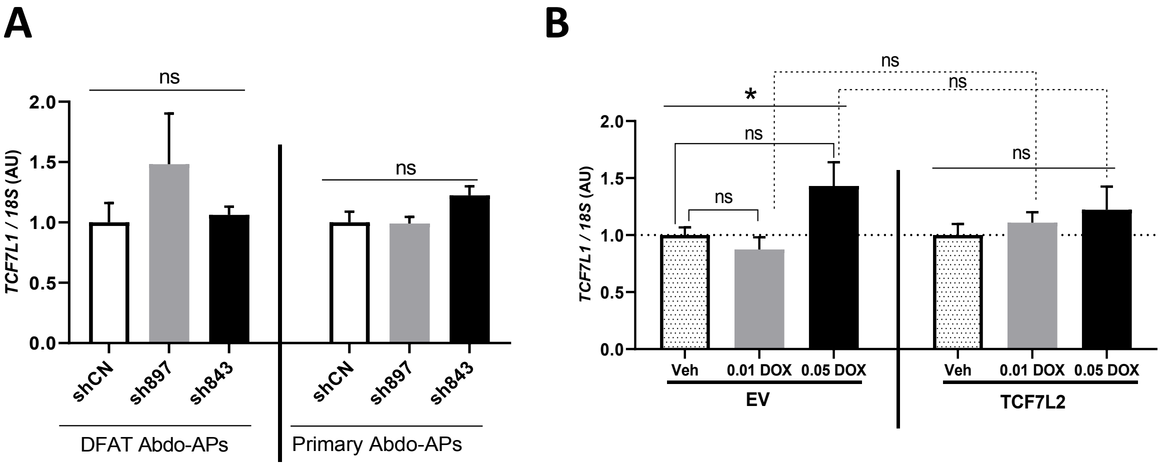
A
ns
ns
TCF7L1 / 18S (AU)
2.0
1.5
1.0
0.5
0.0
shCN
sh897
sh843
shCN
sh897
sh843
DFAT Abdo-APs
Primary Abdo-APs
B
ns
ns
*
ns
ns
ns
TCF7L1 / 18S (AU)
2.0
1.5
1.0
0.5
0.0
Veh
0.01 DOX
0.05 DOX
Veh
0.01 DOX
0.05 DOX
EV
TCF7L2

**Fig. S7**

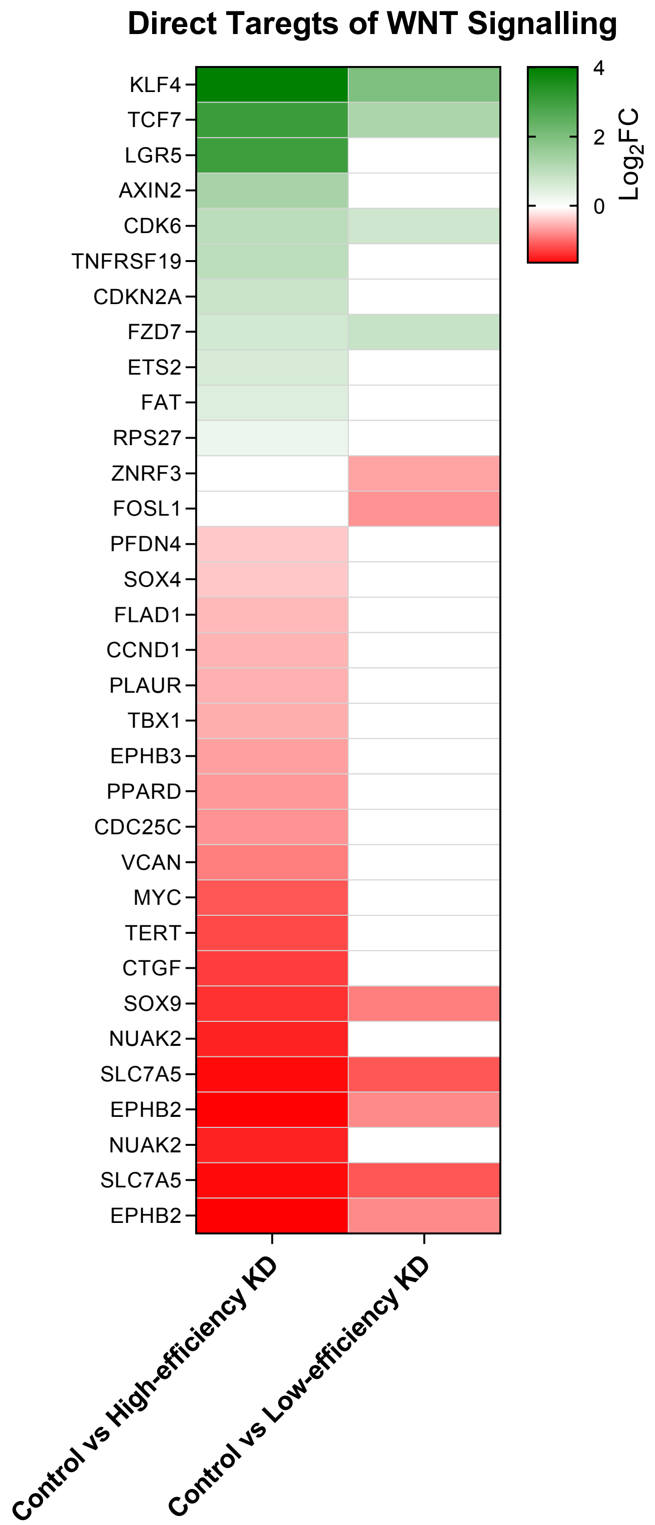
Direct Taregts of WNT Signalling
KLF4
TCF7
LGR5
AXIN2
CDK6
TNFRSF19
CDKN2A
FZD7
ETS2
FAT
RPS27
ZNRF3
FOSL1
PFDN4
SOX4
FLAD1
CCND1
PLAUR
TBX1
EPHB3
PPARD
CDC25C
VCAN
MYC
TERT
CTGF
SOX9
NUAK2
SLC7A5
EPHB2
NUAK2
SLC7A5
EPHB2
4
2
0
$Log_2FC$
Control vs High-efficiency KD
Control vs Low-efficiency KD

### Table. S1

## A

| | BMI (kg/m²) | |
|---|---|---|
| | CC-genotype | TT-genotype |
| N | 600 (336 Female) | 600 (336 Female) |
| Mean | 25.62 | 25.76 |
| SD | 4.157 | 4.261 |
| P25 | 22.67 | 22.8 |
| Median | 25.04 | 25.03 |
| P75 | 27.91 | 27.96 |
| Min | 18.07 | 18.1 |
| Max | 45.96 | 44.75 |

| | Age (yrs) | |
|---|---|---|
| | CC-genotype | TT-genotype |
| N | 600 (336 Female) | 600 (336 Female) |
| Mean | 41.67 | 41.65 |
| SD | 5.67 | 6.01 |
| P25 | 37 | 37 |
| Median | 43 | 43 |
| P75 | 46 | 47 |
| Min | 29 | 29 |
| Max | 53 | 54 |

## B

| CC-genotype | N | Mean | SD | P25 | Median | P75 | Min | Max |
|---|---|---|---|---|---|---|---|---|
| BMI | 3596 (2039F) | 25.85 | 4.6 | 22.6 | 25.15 | 28.24 | 15.29 | 63.14 |
| NEFA | 3594 (2038F) | 485.44 | 236.39 | 316.1 | 445.05 | 612.15 | 29 | 2021.7 |
| Glucose | 3593 (2037F) | 5.18 | 0.45 | 4.87 | 5.15 | 5.45 | 3.28 | 6.99 |
| Insulin | 3596 (2039F) | 13.14 | 6.74 | 9.2 | 11.8 | 15.5 | 0.2 | 99.6 |
| Adipo-IR | 3591 (2036F) | 6359.42 | 4475.95 | 3351.53 | 5227.95 | 8113.05 | 0 | 44712.2 |

| TT-genotype | N | Mean | SD | P25 | Median | P75 | Min | Max |
|---|---|---|---|---|---|---|---|---|
| BMI | 611 (340F) | 25.76 | 4.36 | 22.79 | 25.03 | 27.97 | 17.78 | 46.19 |
| NEFA | 612 (340F) | 477.3 | 217.63 | 315.1 | 457.12 | 620 | 57.6 | 1211 |
| Glucose | 613 (340F) | 5.2 | 0.52 | 4.83 | 5.19 | 5.57 | 3.66 | 6.97 |
| Insulin | 614 (340F) | 12.39 | 6.32 | 8.9 | 11.3 | 14.9 | 1.59 | 97.9 |
| Adipo-IR | 615 (340F) | 5885.88 | 3933.31 | 3135.6 | 4898.5 | 7722.72 | 618 | 31425.9 |
